# Evolutionary rates of testes-expressed genes differ in monogamous and promiscuous *Peromyscus* species

**DOI:** 10.1101/2021.04.21.440792

**Authors:** Landen Gozashti, Russell Corbett-Detig, Scott W. Roy

## Abstract

Reproductive proteins, including those expressed in the testes, are among the fastest evolving proteins across the tree of life. Sexual selection on traits involved in sperm competition is thought to be a primary driver of testes gene evolution and is expected to differ between promiscuous and monogamous species due to intense competition between males to fertilize females in promiscuous lineages and lack thereof in monogamous ones. Here, we employ the rodent genus *Peromyscus* as a model to explore differences in evolutionary rates between testes-expressed genes in monogamous and promiscuous species. We find candidate genes that may be associated with increased sperm production in promiscuous species and gene ontology categories that show patterns of molecular convergence associated with phenotypic convergence in independently evolved monogamous species. Overall, our results highlight the possible molecular consequences of differences in mating system, likely due to differences in selective pressures.

## Main Text

### Introduction

Reproductive genes evolve faster than other genes (Swanson and Vacquier 2002; Clark et al. 2006). This pattern is ubiquitous across diverse lineages, spanning from microbes to mammals (Makalowski and Boguski 1998; Armbrust and Galindo 2001; Sato et al. 2002; Torgerson et al. 2002; Wik et al. 2008; Vacquier and Swanson 2011). Testes-expressed genes in particular are fast evolving, likely due to sexual selection driven by sperm competition (Parker 1970; Civetta and Singh 1995; Rice and Holland 1997; Swanson and Vacquier 2002; Torgerson et al. 2002; Ramm and Stockley 2009; Firman and Simmons 2010a; Harrison et al. 2015; Moyle et al. 2020). Rapid evolution of testes-expressed genes is thought to generally reflect positive selection (Ramm et al. 2008; Ramm and Stockley 2009; Teng et al. 2017), but could also be explained by relaxed selection (Dapper and Wade 2016).

Differences in mating system, that is the general pattern by which males and females mate within a species, are expected to drive differences in rates of evolution of testes-expressed genes (Lüpold et al. 2016; Clutton-Brock 2017). Most mammalian species are promiscuous, meaning that females mate with multiple males and vice versa (Garcia-Gonzalez 2017). However, genetic monogamy (where males and females only mate with one individual for life) has evolved multiple times independently across mammalian lineages (Lukas and Clutton-Brock 2013). Promiscuous and monogamous species often exhibit morphological, physiological and behavioral differences associated with mate fidelity (Stanyon and Bigoni 2014; Tidière et al. 2015; Dapper and Wade 2016; Wey et al. 2017; Ambaryan et al. 2019; Civetta and Ranz 2019). These differences are often associated with the male testes and with sperm performance, consistent with promiscuous but not monogamous males competing to fertilize females (Firman and Simmons 2010a; Dapper and Wade 2016). Faster, more viable sperm and higher sperm counts yield an increased chance of fertilization in promiscuous species (Pizzari 2006). Indeed, promiscuous males often possess larger testes and produce more abundant and more competitive sperm than monogamous males (Heske and Ostfeld 1990; Shuster 2009; Firman and Simmons 2010a; Claw et al. 2018; Fisher et al. 2018). However, the genetic basis of morphological and physiological differences related to differences in mating system remain largely unexplored.

Social/sexual monogamy has evolved at least twice independently within the rodent genus *Peromyscus* (Figure 1) (Turner et al. 2010; Bedford and Hoekstra 2015), with behavioral and physical traits differing consistently between monogamous and promiscuous *Peromyscus* species (Fisher and Hoekstra 2010; Fisher et al. 2014, 2016, 2018; Bendesky et al. 2017). *Peromyscus* is a powerful model for interrogating the drivers of convergent evolution and adaptation due to its well-resolved phylogeny (Greenbaum et al. 2017; (Sullivan et al. 2017), convergence of phenotypes (Steiner et al. 2009; Manceau et al. 2010; Bedford and Hoekstra 2015), and ecological, morphological, and behavioral variation (Gering et al. 2009; Fisher and Hoekstra 2010; Shorter et al. 2012; Bedford and Hoekstra 2015; Guralnick et al. 2020). Here, we combine evolutionary rate modeling and gene expression analysis to identify genes and gene ontology categories associated with differential evolution in promiscuous and monogamous species.

**Figure 1:**
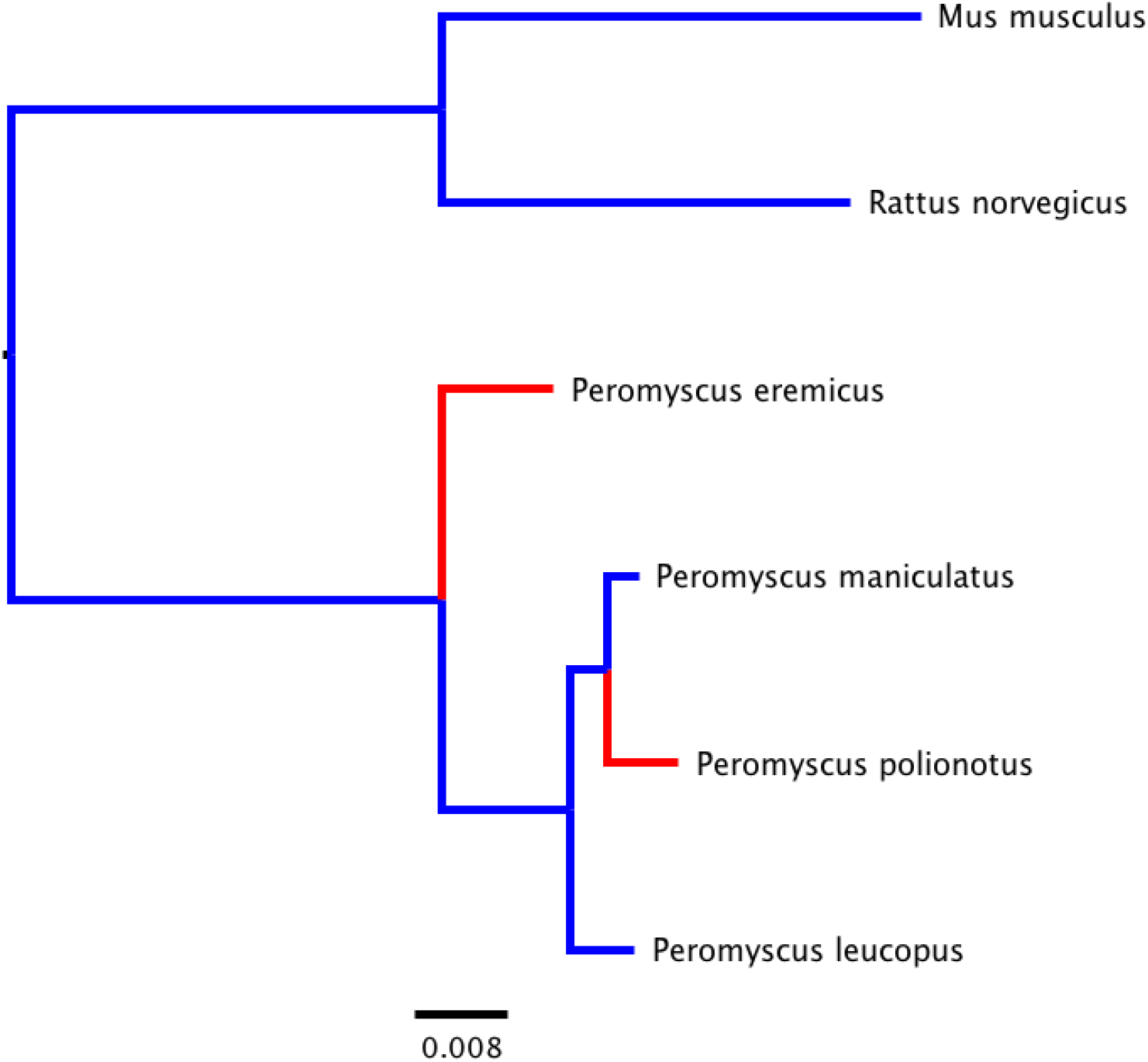
Phylogenetic relationships of four considered *Peromyscus* species with the addition of *Mus musculus* and *Rattus norvegicus* as outgroups. Our tree was produced with *IQ-TREE* (Minh et al. 2020) using 15 concatenated orthologs in each respective species. Genetic distances are measured in units of the number of base substitutions in proportion to alignment length. Blue colored nodes represent promiscuity and red colored nodes represent monogamy

## Results and Discussion

Maximum-likelihood analysis reveals 6 genes exhibiting significant differences in evolutionary rates between the monogamous *P. eremicus* and the promiscuous *P. maniculatus*. Intriguingly, 5 of these are implicated in spermatogenesis and are evolving faster in *P. maniculatus*: Znf644, KDM3A, Ddx25, Spata16 and ESSBP (Figure 2, Table 1) (Dam et al. 2007; Liu et al. 2010; Tsai-Morris et al. 2010; Souza et al. 2017; Wilson et al. 2017). Sequence level changes leading to increased sperm abundance or competitive ability might be favored by selection in promiscuous species due to the selective pressures of mate competition. Thus, we hypothesize that these genes may have experienced positive selection in *P. maniculatus*. Previous studies have suggested that sperm production is sensitive to local adaptation in light of the pressures associated with competition (Ramm and Stockley 2009; Winkler et al. 2019; Lindsey et al. 2020).

**Table 1:** Genes exhibiting significantly different rates of evolution in considered monogamous and promiscuous species. Column 1 displays gene name. Column 2 provides the trend of evolution (monogamous (M) species vs. promiscuous (P) species). Column 3 summarizes respective known involvement in reproduction, column 4 provides FDR corrected X^2^ P-values (Q-values), and column 5 contains sources for sexual implications.

| Gene | Relative Evolutionary Rate | Involvement in Reproduction | Q value | Source for sexual implication |
| --- | --- | --- | --- | --- |
| KDM3A | P>M | Spermatogenesis | <0.0001 | (Wilson et al. 2017) |
| Spata16 | P>M | Spermatogenesis | 0.0006 | (Chang et al. 2004) |
| Ddx25 | P>M | Spermatogenesis | 0.0020 | (Tsai-Morris et al. 2010) |
| Znf644 | P>M | Spermatogenesis | 0.0002 | (Liu et al. 2010) |
| ESSBP | P>M | Spermatogenesis | <0.0001 | (Souza et al. 2017) |
| Arhgef18 | P>M | Cell morphology and cell-cell interactions | <0.0001 | (Niu et al. 2003) |

**Figure 2:**
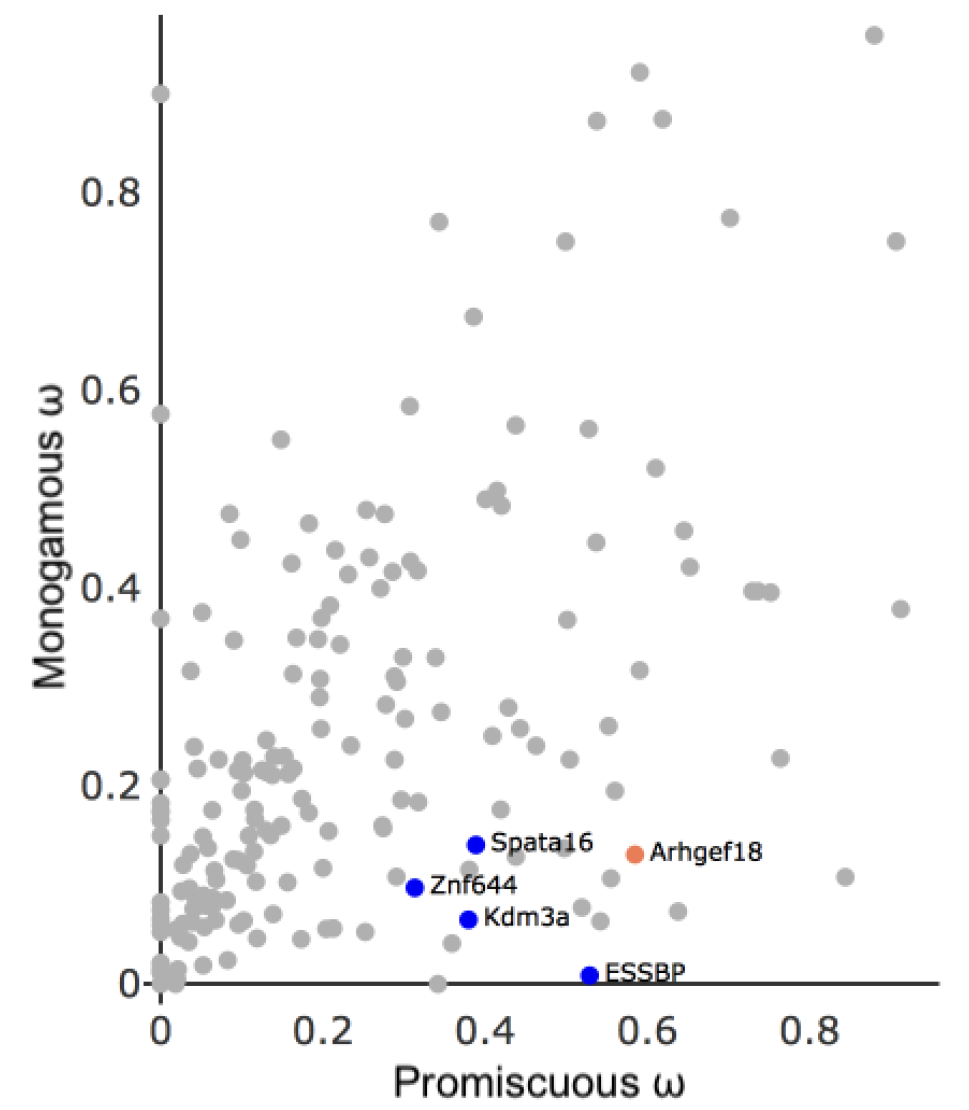
Respective evolutionary rates (ω) for genes exhibiting different evolutionary rates between in the monogamous *P. eremicus* and the promiscuous *P. maniculatus*. Colored points correspond to genes exhibiting statistically significant differences in evolutionary rates. Blue points correspond to genes that play crucial roles in spermatogenesis.

Arhgef18, the other gene with a significantly higher evolutionary rate in the promiscuous *P. maniculatus*, lacks known reproductive implications. Although its role in spermatogenesis remains unexplored, Arhgef18 plays a role in maintaining apico-basal polarity and localization of tight junctions and cortical actin. Cellular morphology and cell-cell interactions are essential to sperm cell organization and sertoli-spermatid interactivity during spermatogenesis (Wong and Cheng 2009; Gao and Cheng 2016). Its rapid evolution in *P. maniculatus* suggests it too may evolve under positive selection.

Differential gene expression analysis lends further support for positive selection of Spata16 and ESSBP. We calculated change in RPKMs for homologous transcripts between species, focusing specifically on the 6 genes exhibiting significantly different rates of evolution. Interestingly, we find that ESSBP and Spata16 exhibit significantly increased expression in *P. maniculatus* (*Mann-Whitney U* P = 0.0294 and 0.0376 respectively). The relationship between sequence divergence and expression divergence between species has been shown to be positively correlated in mammals (Liao and Zhang 2006; Warnefors and Kaessmann 2013). This pattern is consistent with changes in gene expression levels resulting from positive selection (Jordan et al. 2005), further raising the possibility that ESSBP and Spata16 might be under positive selection in *P. maniculatus*.

We also used branch models to compare shared rates of evolution between two species in which monogamy evolved independently, *P. eremicus* and *P. polionotus* and two relatively divergent promiscuous species, *P. maniculatus* and *P. leucopus*. Due to the recent divergence of *P. maniculatus* and *P. polionotus*, branch tests using individual genes may be unreliable. Therefore, rather than focusing on significantly different rates between independent homologous loci, we used MGI’s gene ontology database (Smith and Eppig 2009) to cluster all considered genes based on ontology, and explored trends of shared evolutionary rates between genes in monogamous species relative to promiscuous species. We find significant differences in evolutionary trends between monogamous and promiscuous species in eight gene ontology categories, five of which are important for sperm function (Table 2, Figure 3). We find that genes within ontologies with implications in sperm function generally have slightly increased evolutionary rates in monogamous species. In fact, all considered genes involved in “flagellated sperm motility,” “motile cilium,” “centrosome” and “cell projection” are evolving at faster rates in monogamous species. One possible explanation for this is that monogamous species exhibit reduced selective constraint on genes integral to sperm function. Mutations are likely filtered more frequently by purifying selection in promiscuous species than in monogamous species where negative selection is relaxed (Firman and Simmons 2010a). An alternative explanation is positive selection on these genes in monogamous species. However, in contrast with the expected observation for loci under positive selection, we do not find significant differences in evolutionary rates between monogamous and promiscuous species when we compare genes within these ontologies independently, and instead find that they exhibit dN/dS values closer to 1. These observations may constitute a molecular signature of convergent evolution in protein evolutionary rates for genes involved in sperm competitiveness in monogamous species.

**Table 2:** MGI Ontology categories with differences in trends of evolutionary rates.

| MGI Ontology | Relative Evolutionary Rate | Sexual Implication | P value | Source for sexual implication |
| --- | --- | --- | --- | --- |
| cell projection | M>P | Sperm function (motility) | 0.0078 | (Avidor-Reiss et al. 2017) |
| centrosome | M>P | Sperm function (motility) | 0.0078 | (Terada et al. 2010; Schatten et al. 2011; Schatten and Stearns 2015; Gunes et al. 2020) |
| cilium | M>P | Sperm function (motility) | 0.0390 | (Gadella 2008; Inaba and Mizuno 2016; Yuan et al. 2019; Sironen et al. 2020) |
| flagellated sperm motility | M>P | Sperm function (motility) | 0.0156 | (Vladić et al. 2002; Lüpold et al. 2009; Firman and Simmons 2010b; Fisher et al. 2016) |
| hydrolase activity | M>P | NA | 0.0351 | NA |
| motile cilium | M>P | Sperm function (motility) | 0.0312 | (Gadella 2008; Inaba and Mizuno 2016; Yuan et al. 2019; Sironen et al. |
|  |  |  |  | 2020) |
| nucleic acid binding | M>P | NA | 0.0352 | NA |
| ribonucleoprotein complex | M>P | NA | 0.0313 | NA |

**Figure 3:**
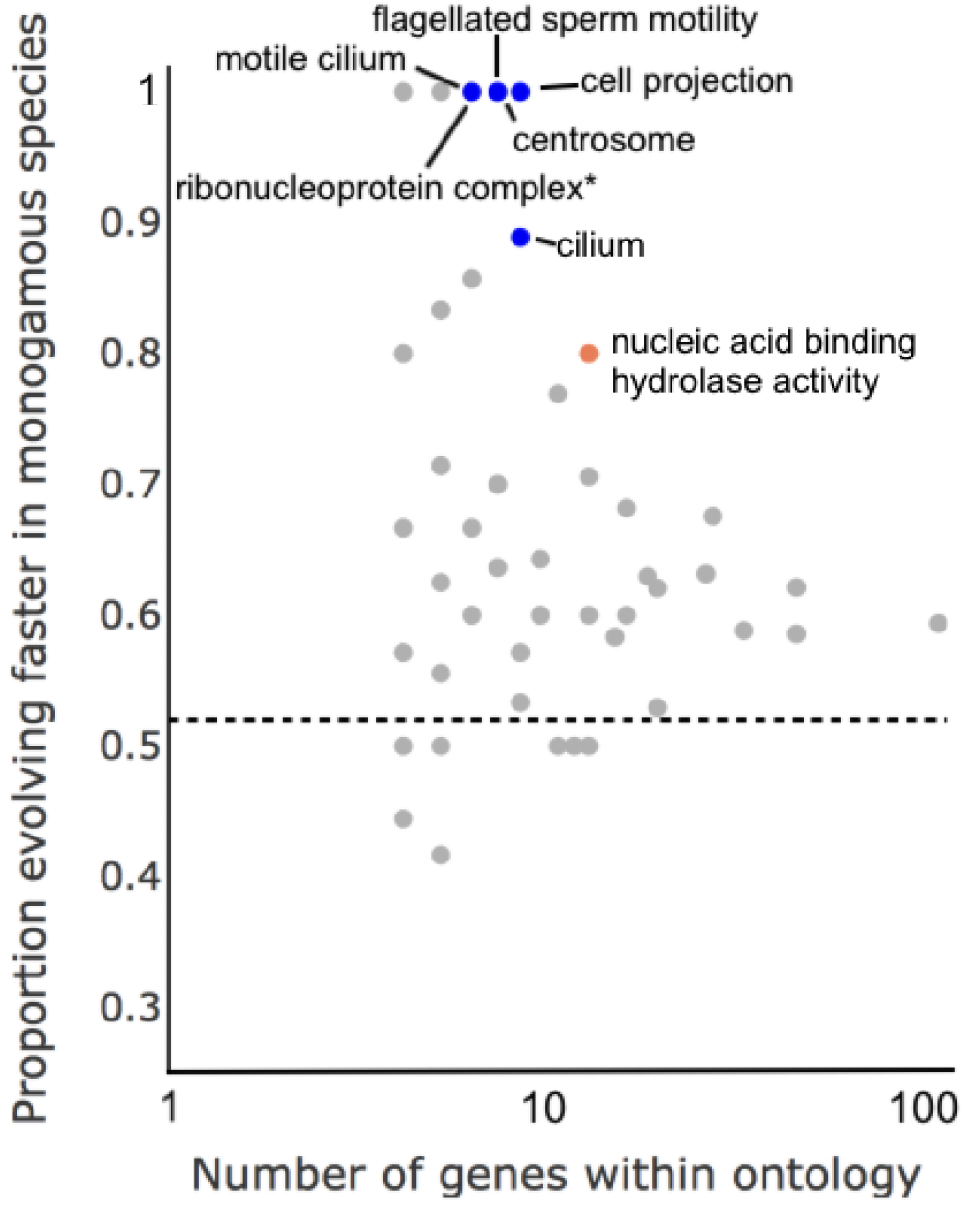
Proportion of genes evolving faster in monogamous species relative to the number of genes within each respective ontology cluster. The five Ontology groups with significant trends of evolution in monogamous species relative to promiscuous ones and known specific implications in sperm performance are colored blue. Other significant ontologies are colored in orange. The dotted line represents the null expected proportion of genes evolving faster in monogamous species. The “*” denotes an ontology group that is not specifically involved in sperm performance but whose data point overlaps with an ontology group. We note that for visualization purposes, we only show gene ontology groups containing four or more genes, and that there is an aggregation of smaller ontology groups not shown which account for the apparent skew of points above the expected proportion of genes evolving faster in monogamous species.

Our results illuminate the molecular correlates of changes in reproductive functions associated with changes in mating systems. Sperm abundance, motility and morphology are critical for fertility (Sharpe 2012; Baptissart et al. 2013). We identify six genes that exhibit significantly increased evolutionary rates in the promiscuous *P. maniculatus* relative to the monogamous *P. eremicus*. Five of these are known to be involved in spermatogenesis and the sixth has plausible connections. Additionally, two genes exhibit significantly increased expression in *P. maniculatus*, providing additional evidence that these loci experience positive selection in *P. maniculatus* due to competition between males to fertilize females. Previous studies have found that sperm in the promiscuous *P. maniculatus* exhibit adaptive morphological traits and increased abundance when compared to that of monogamous *Peromyscus* species (Fisher et al. 2016).

Furthermore, genes implicated in sperm function exhibit marginally faster evolutionary rates in two independently evolved monogamous species relative to promiscuous ones when clustered by ontology, highlighting the possible role of relaxed selection. However, future work is required to investigate roles of selection, and we propose that these genes serve as candidates for future studies on adaptation and convergent evolution with regard to reproductive behavior. Future studies should include nucleotide and transcript-abundance polymorphism data to enable direct estimation of the mode of selection and increased sampling of monogamous species across many independent origins in diverse lineages to test for molecular convergence associated with differences in mating system.

## Materials and Methods

### Obtaining and reconstructing transcriptomic data

We developed a pipeline to detect signatures of parallel evolutionary rates in transcripts of *P. polionotus* and *P. eremicus* relative to *P. leucopus* and *P. maniculatus*. First, we obtained *P. maniculatus, P. polionotus, P. leucopus*, and *P. eremicus* testes transcriptome data from (Lindsey et al. 2020), (https://www.ncbi.nlm.nih.gov/bioproject/584485), (Harris et al. 2013) and (Kordonowy and MacManes 2016) respectively. The testes specific transcriptome for *P. eremicus* has previously been assembled (Kordonowy and MacManes 2016). For the other three species, we aligned each species’ testes transcriptome data to reference CDS data (GCF_000500345.1 for *P. maniculatus* and *P. polionotus* since no reference exists for *P. polionotus* and *P. maniculatus* is its closest relative with a reference CDS file, and GCF_004664715.2 for *P. leucopus*) using *hisat2 (Kim et al. 2015)* and called variants using *bcftools (Danecek et al. 2021)*. We then reconstructed each transcript using a custom python script (see https://github.com/lgozasht/Peromyscus-reproductive-genetic-differences).

### Identification of orthologous gene sets

We performed an all vs. all BLAST (Altschul et al. 1990) with a minimum 90 percent identity and 80 percent sequence overlap to identify putative homologous transcripts between species. Then, we employed *silixx* (Miele et al. 2011) to cluster transcripts into families of highly similar sequences from homologous genes. We filtered for families that contained transcripts from all four species and included the transcript with the lowest e-value when a species possessed multiple candidate transcripts for a given family. We identified 942 transcripts that shared homology between all 4 of our considered species.

### Generating and curating PAML input files

For each transcript, we used *MAFFT (Katoh and Standley 2013)* to generate a multiple sequence alignment and *IQ-TREE* (Minh et al. 2020) to infer a tree using maximum likelihood. Since indels and poor alignment quality can impair *PAML* functionality, we performed additional filtering on our alignments. We used *Alignment_Refiner_v2, (Young et al. 2019)* to trim alignments above an overall missing data threshold of 1% and removed clusters with greater than 50 gaps in a particular sequence or 100 total gaps. We also removed alignments containing sequences with lengths not divisible by 3. We were left with 199 transcripts suitable for PAML input after filtering

### Comparing evolutionary rates of transcripts in P. eremicus and P. maniculatus

We employed *PAML* (Yang 2007) to search for shared signatures of evolutionary rates in *P. eremicus* and *P. polionotus* relative to *P. leucopus* and *P. maniculatus*. Short branch lengths between *P. maniculatus* and *P. polionotus* could inhibit accurate pairwise *dN/dS* comparisons (Yang 2007; Kryazhimskiy and Plotkin 2008; Mugal et al. 2014), so instead we performed comparisons between *P. maniculatus* and *P. eremicus*. For each transcript cluster, we applied *PAML* with a null model (*model = 2*) in which *dN/dS* differs in *P. maniculatus* and *P. eremicus* relative to *P. polionotus* and *P. leucopus*, and with an alternative model (*model = 2*) in which each *dN/dS* differs between *P. maniculatus* and *P. eremicus* relative to *P. polionotus* and *P. leucopus*. To assess significance, we used a likelihood ratio test, which is X^2^ distributed with a single degree of freedom. After further manual alignment filtering, we applied a false-discovery rate correction for multiple testing (Benjamini and Hochberg 1995) yielding 6 significant transcripts for downstream analysis (α = .05).

### Differential expression analysis of P. eremicus and P. maniculatus

We obtained testes RNA data from three runs of separate replicates for *P. eremicus* (SRA run excessions: ERR1353571, ERR1353572, and ERR1353573) and one run of pooled replicates for *P. maniculatus* (SRA run excession: SRR8587279) and aligned to respective testes transcriptomes using *hisat2* (Kim et al. 2015). Next, we obtained read counts for each transcript using *htseq-count* (Anders et al. 2014) and calculated *RPKM* for each transcript. We performed independent Mann–Whitney U tests for the change in *RPKMs* of transcripts corresponding to each gene in which we observed significant differences in evolutionary rates between *P. eremicus* and *P. maniculatus*. We aggregated p-values for transcripts corresponding to each gene using Fisher’s combined probability to obtain a final p-value for each gene (Yi et al. 2018).

### Comparing parallel rates of evolution between monogamous and promiscuous species

We used *PAML* to compute the likelihood of a null model (*model = 0*) in which *dn/ds* is equal across all branches of the tree with our alternative model. Then we fit a branch model (*model = 2* in which the *dn/ds* ratio is allowed to differ between monogamous and promiscuous lineages, but where the two ratios are constrained to be identical across all monogamous or promiscuous branches. We again used a likelihood ratio test to assess significance.

### Finding evolutionary trends in different gene ontologies

We interrogated trends of evolutionary rates by aligning each gene that passed our upstream filtering to the *Mus musculus* GRCm38 genome (Genbank accession GCF_000001635.26) and cross referencing the MGI ontology database (see http://www.informatics.jax.org/mgihome/projects/aboutmgi.shtml). For each group of genes in a given GO term, we performed a two-sided binomial test with regard to the number of genes evolving at slower (or faster) rates in monogamous species relative to promiscuous species. For our null, we used the proportion of genes evolving at slower rates between monogamous and promiscuous species among all considered loci: P = 0.519. We filtered GO groups with significant binomial P-values (*α* < 0.05). To determine that these signatures of correlated evolutionary rates within GO categories are not driven by species-specific effects, we tested for idiosyncratic evolution in each species. To do this, we ran *PAML* with a branch model in which each species evolved at a unique rate relative to the other three.

## Acknowledgments

The authors thank Jenny Chen for her input on optimizing statistical analyses and Hopi Hoekstra for advice. The authors declare no conflicts of interest. This research did not receive any specific grant from funding agencies in the public, commercial, or not-for-profit sectors.

